# Signal-regulated unmasking of the nuclear localization motif in the PAS domain regulates the nuclear translocation of PASK

**DOI:** 10.1101/2023.09.06.556462

**Authors:** Michael Xiao, Sajina Dhungel, Roksana Azad, Denize C. Favaro, Rajaian Pushpabai Rajesh, Kevin H. Gardner, Chintan K. Kikani

**Affiliations:** Department of Biology, University of Kentucky, Lexington, KY 40502, USA; Structural Biology Initiative, CUNY Advanced Science Research Center, New York, NY 10031, USA; Ph.D. Program in Biochemistry, Graduate Center, City University of New York, NY 10016, USA; Department of Chemistry and Biochemistry, City College of New York, NY 10031, USA; Ph.D. Programs in Biochemistry, Chemistry and Biology Ph.D. Programs, Graduate Center, City University of New York, NY 10016, USA

## Abstract

The ligand-regulated PAS domains are one of the most diverse signal-integrating domains found in proteins from prokaryotes to humans. By biochemically connecting cellular processes with their environment, PAS domains facilitate an appropriate cellular response. PAS domain-containing Kinase (PASK) is an evolutionarily conserved protein kinase that plays important signaling roles in mammalian stem cells to establish stem cell fate. We have shown that the nuclear translocation of PASK is stimulated by differentiation signaling cues in muscle stem cells. However, the mechanistic basis of the regulation of PASK nucleo-cytoplasmic translocation remains unknown. Here, we show that the PAS-A domain of PASK contains a putative monopartite nuclear localization sequence (NLS) motif. This NLS is inhibited in cells via intramolecular association with a short linear motif, termed the PAS Interacting Motif (PIM), found upstream of the kinase domain. The interaction between the PAS-A domain and PIM is evolutionarily conserved and serves to retain PASK in the cytosol in the absence of signaling cues. Consistent with that, we show that metabolic inputs induce PASK nuclear import, likely by disrupting the PAS-A: PIM association. We suggest that a route for such linkage may occur through the PAS-A ligand binding cavity. We show that PIM recruitment and artificial ligand binding to the PAS-A domain occur at neighboring locations that could facilitate metabolic control of the PAS-PIM interaction. Thus, the PAS-A domain of PASK integrates metabolic signaling cues for nuclear translocation and could be targeted to control the balance between self-renewal and differentiation in stem cells.

## Introduction

The ligand-regulated PAS domains are one of the most diverse signal-integrating domains found in proteins from prokaryotes to humans [1, 2], linking organic and inorganic environmental cues to cellular signaling via various cognate functional domains [3, 4]. Thus, PAS domains facilitate dynamic cellular responses by biochemically connecting cellular processes with changing environmental landscapes.

PAS domain-containing Kinase (PASK) is an evolutionarily conserved protein kinase that combines PAS sensory domains with a catalytic CAMK-type Ser/Thr kinase domain [5, 6]. PASK performs important signaling functions in mammalian stem cells to establish stem cell fate [6, 7]. PASK is highly expressed in proliferating, undifferentiated embryonic and adult stem cells of various cell lineages but declines as stem cells differentiate, ultimately being nearly completely lost in differentiated cells and tissues [7]. As a sensory kinase enriched in stem cells, PASK plays a critical role in balancing the self-renewal or differentiation pathways in accordance with the metabolic environment in stem cells. In proliferating, self-renewing stem cells, PASK predominantly resides in the cytosol; however, it is translocated to the nucleus in response to differentiation signaling cues, with this PASK nuclear translocation being essential for the disruption of a mitotic self-renewal network [8]. We showed that the nuclear translocation of PASK is signal-regulated, stimulated by mitochondrial glutamine metabolism [8]. Glutamine depletion retained PASK in the cytoplasm, strengthening the mitotic self-renewal network and inhibiting the differentiation program. Thus, understanding the mechanism of nucleo-cytoplasmic distribution of PASK is key to understanding how signaling pathways establish differentiation competence.

Here we elucidate the mechanistic basis for the signal-regulated control of PASK nuclear translocation. In our current study, we unexpectedly identified a novel intramolecular interaction involving the PASK N-terminal PAS domain (PAS-A) and a short-linear PAS interacting motif (PIM) in the C-terminal region of PASK that masks a signal-regulated nuclear localization motif within PAS-A. The PAS-A:PIM interaction occurs adjacent to a cavity within PAS-A where we have previously demonstrated small molecule binding, suggesting a potential novel allosteric control of intramolecular association in PASK. Taken together, our data provide avenues for precise metabolic control of PASK subcellular distribution and catalytic activity.

## Results

### Identification and characterization of the PAS-A interacting motif in PASK *in vivo*

In proliferating mammalian stem cells, full-length WT PASK is retained in the cytoplasm [8]. At the onset of differentiation, a large proportion of PASK is translocated to the nucleus to catalytically activate the terminal differentiation program [8]. Therefore, we hypothesized that PASK contains a signal-regulated mechanism for the temporal regulation of PASK nuclear translocation. To evaluate this hypothesis, we set out to first identify sequence features in PASK that mediate the nuclear import of PASK, which could be targeted by signaling pathways during myogenesis. We performed a domain truncation analysis to identify regions in PASK that play important roles in nucleo-cytoplasmic trafficking. Human PASK is 1323 amino acids long and is predicted to contain three clearly identifiable domains: two PAS domains, including PAS-A, (experimentally confirmed between AA 131-237) [9] and PAS-B (predicted only, ca. AA 254-421) [6], and a C-terminal CAMK-type Ser/Thr kinase domain (experimentally confirmed between AA 977-1300) [5] separated from the two PAS domain by a ∼600 residue-long unstructured region. Considering these structural landmarks, we generated a series of truncations (Figure 1A, Table 1) in PASK and analyzed their cellular localizations in HEK293T cells (Figure 1B). In contrast to full-length WT PASK [8], we noticed that a GFP-tagged C-terminal fragment (Fragment A, AA 841-1323) was localized into the nucleus to varying degrees when expressed individually. In contrast, the N-terminal PAS^737^ fragment (AA 1-737, including PAS-A) was predominantly localized in the cytosol in HEK293T cells when expressed alone (Figure 1B), likely due to the presence of the recently-identified Nuclear Export Sequences (NES) within its sequence [8].

**Figure 1:**
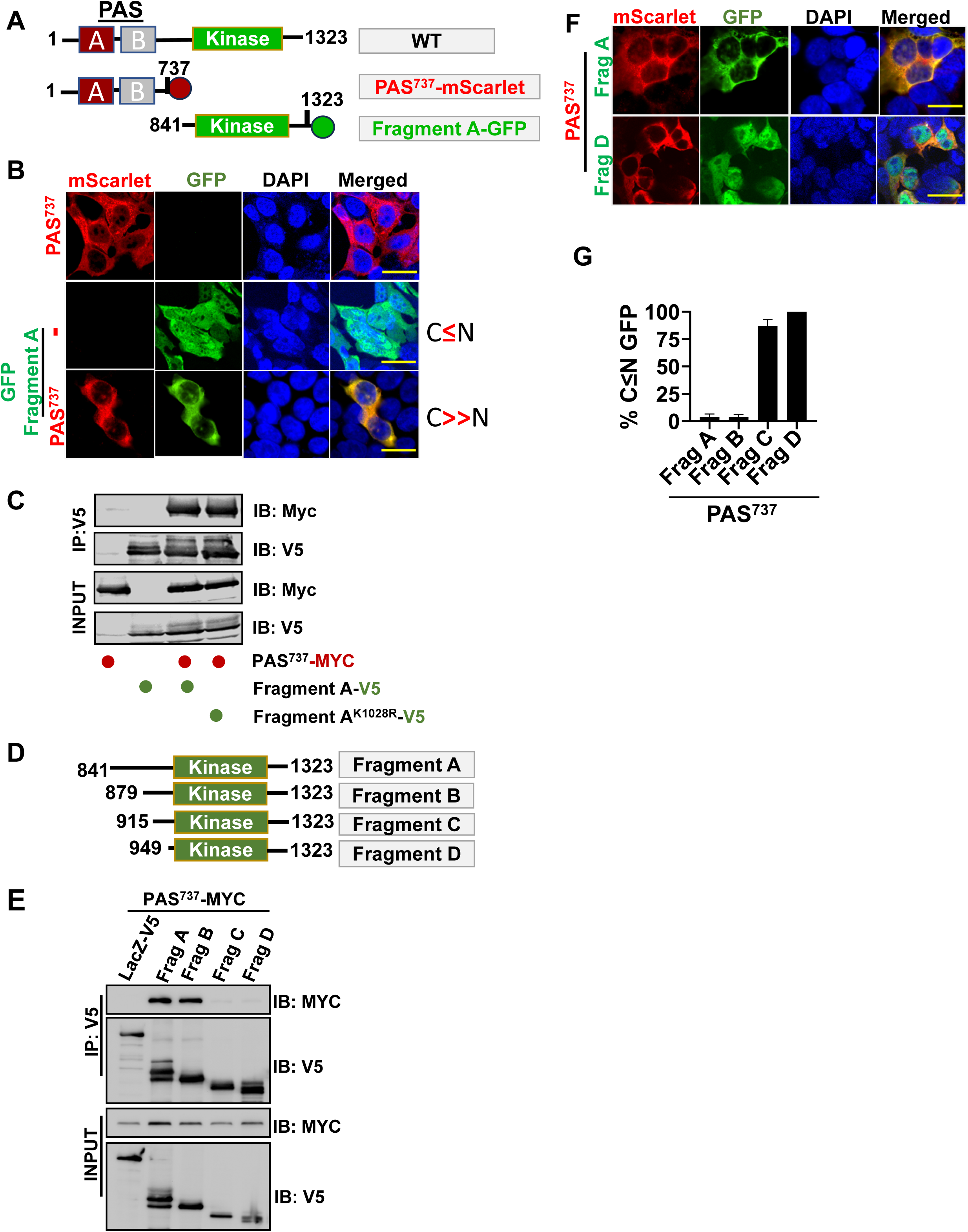
Identification of intramolecular interactions in PASK as regulators of PASK nuclear localization. **(A)** Domain architecture of human PASK, including two PAS domains (the experimentally verified PAS-A (red) and the PFAM-predicted PAS-B (gray), and the serine/threonine kinase catalytic domain (green)). The PAS^737^ fragment consists of amino-acids 1-737 and lacks the kinase domain, while Fragment A consists of amino acids 841-1323 and lacks both PAS domains. **(B)** GFP-tagged Fragment A was expressed in HEK293T cells alone, or together with mScarlet-tagged PAS^737^. 24 hr after transfections, cells were fixed and relative distribution of Fragment A and PAS^737^ was determined by confocal microscopy. Scale bar = 40 µM. **(C)** WT or kinase-dead (K1028R) version of V5-tagged Fragment A were expressed alone or together with Myc-tagged PAS^737^ in HEK293T cells, and their interaction was determined by immunoprecipitation. **(D)** Domain architecture of C-terminal truncations (Fragments A to D) generated to identify region of PASK that interacts with N-terminal PAS^737^ region. **(E)** Various V5-tagged C-terminal fragments were co-expressed with PAS^737^-Myc and co-immunoprecipitation was conducted as described in Materials and Methods. **(F)** GFP-tagged Fragments A and D were expressed in HEK293T cells along with mScarlet-tagged PAS^737^ in HEK293T cells. 24hr after transfections, cells were fixed and relative distribution of fragments (A and D) and PAS^737^ was determined by confocal microscopy. Scale bar = 40 µM. **(G)** Quantification of % cells with nuclear-localized GFP from experiment in Figure 1F.

Intriguingly, co-expression of PASK Fragment A (C-terminal region AA 841-1323) with PAS^737^ (N-terminal region AA 1-737) resulted in the nuclear exclusion of Fragment A (Figure 1B), raising the possibility of an association between PAS^737^ and Fragment A, leading to the retention of Fragment A in the cytosol. Indeed, PAS^737^ and Fragment A associated in co-immunoprecipitation experiments in HEK293T cells (Figure 1C). Since Fragment A contains a kinase domain, we also asked if PASK catalytic activity was required for binding PAS^737^; mutation of a key ATP-binding residue (K1028R) did not affect the PAS^737^: Fragment A co-immunoprecipitation, suggesting that the autophosphorylation activity of PASK is not involved in regulating this association.

To map the PAS^737^-interacting residues within Fragment A, we performed a series of truncation analyses from the N-terminal end of Fragment A (Figure 1D). As shown in Figure 1E, we noticed strong interactions between PAS^737^ and Fragments A (AA 841-1323) and B (AA 879-1323), but not with the shorter Fragments C (AA 915-1323) and D (AA 949-1323). Thus, our data suggested that residues between 879 and 915 interact with the N-terminal PAS^737^ Fragment. Consistent with the hypothesis that the PAS domain affects the subcellular distribution of C-terminal fragments by physical association, we noticed that Fragments A and B were retained in the cytosol when co-expressed with the PAS^737^ fragment in HEK293T and C2C12 cells (Figures 1F, G; Figure S1). In contrast, Fragments C and D, which did not interact with the PAS^737^ fragment, remained nuclear in PAS^737^ expressing cells in C2C12 cells (Figure S1). These results suggest that an intramolecular association in PASK, here emulated by the intermolecular binding of PAS^737^ with the C-terminal Fragments A and B, might dynamically regulate its nucleo-cytoplasmic shuttling.

Because PASK is an evolutionarily conserved protein kinase, we reasoned that the functionally important residues for Fragment A (841-1323) and PAS^737^ Fragment interaction would likely be conserved. Indeed, multiple sequence alignment of PASK across various species showed strong sequence conservation between amino acids 879 and 915 (Figure 2A). We particularly noticed the presence of a GXXXG-like motif (G^892^-A^893^-Y^894^-S^895^-G^896^) and the highly conserved Y^894^ and S^897^ residues, which could be targeted by posttranslational modifications. Mutations of either the GXXXG motif with AXXXA (Figure 2B, Table 1, designated as PIM-Mut1) or Y894A and S897A (Table 1, Figure 2B, Table 1, designated as PIM-Mut2) substantially disrupted the co-immunoprecipitations between the mutated Fragment A with PAS^737^, indicating that these residues are necessary for the intramolecular interaction between PAS^737^ and Fragment A. Furthermore, a Flag-GFP fusion construct with a peptide comprising the minimal conserved region (Table 1, AA 879-904, termed PIMtide^WT^) was sufficient to interact with PAS^737^ (Figure 2C, D); whereas the Y894AS897A mutated PIMtide (Table 1, AA 879-904 (Y894AS897A), termed PIMtide^Mut2^) completely disrupted this association, just as it did in the larger Fragment A context. Functionally, while WT-Fragment A was retained in the cytosol when co-expressed with PAS^737^, the Y894AS897A mutated Fragment A remained nuclear and unaffected by co-expression of PAS^737^ fragment (Figures 2E, F). Thus, residues 879-904, which we term <u>P</u>AS Interacting <u>M</u>otif, or PIM, are necessary and sufficient to mediate an intermolecular interaction between N-terminal and C-terminal regions within PASK that we suspect to also work in an intramolecular context.

**Figure 2:**
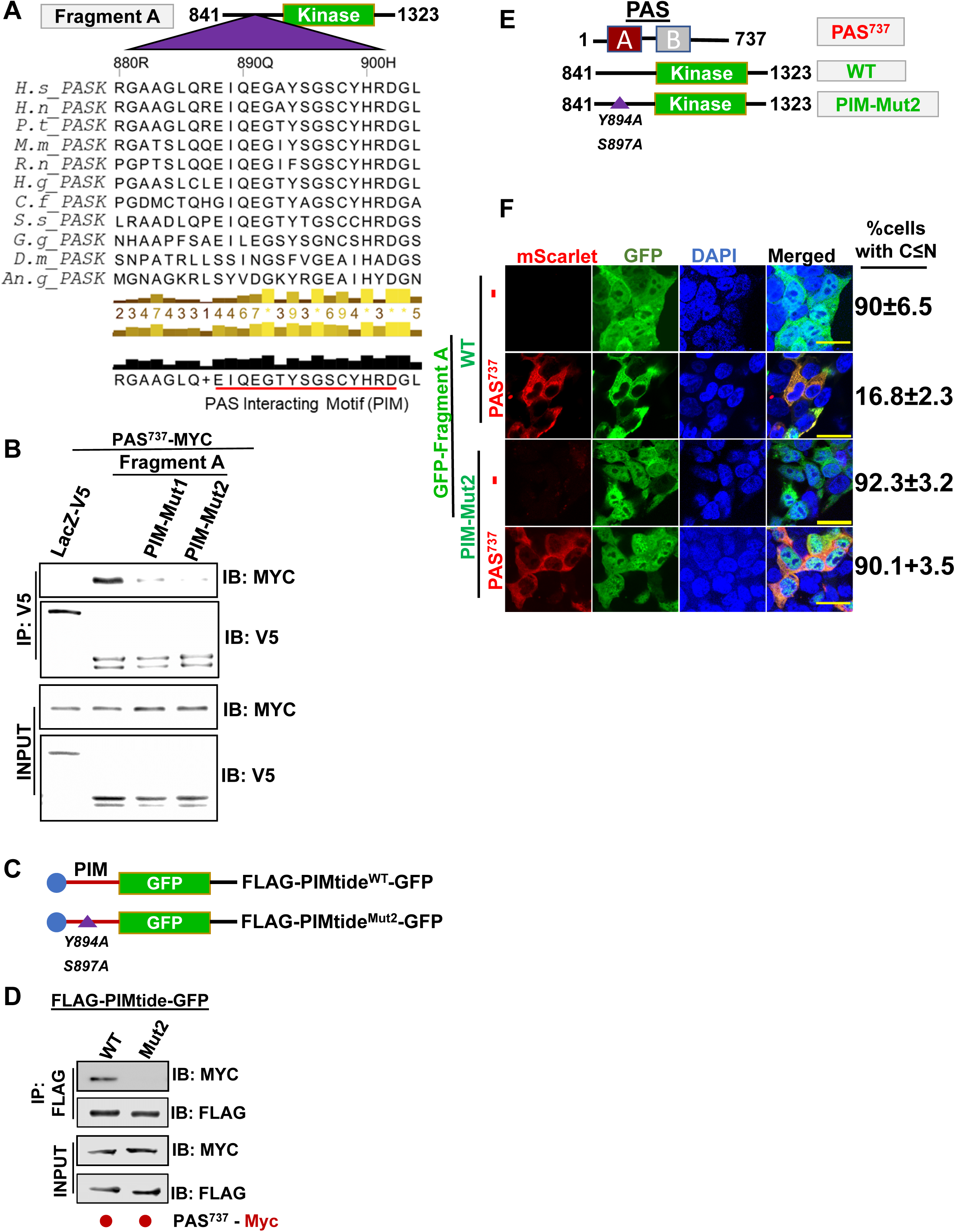
Identification of residues Y894 and S897 as critical for intramolecular interactions in PASK. **(A)** Primary sequence alignment of PASK from invertebrates and vertebrates. The PAS Interacting Motif (PIM) sequence is depicted by a red line. The yellow and black bars at the bottom of the alignment indicate positional similarity scores. **(B)** Fragment A mutants, G891A/G895A (PIM-Mut1) or Y894A/S897A (PIM-Mut2) were co-expressed with PAS^737^-Myc in HEK293T cells. Co-IP was performed as described in methods and material. **(C-D)** WT or Mut2 version of PIM sequence (PASK AA 879-904) was fused with Flag-GFP (PIMtide). The sufficiency of PIM interaction with PAS^737^ region was evaluated by co-immunoprecipitation of HEK293T co-expressed proteins. **(E)** Schematic of PAS^737^ and Fragment A-WT or Mut2 used in Figure 2F. **(F)** Nuclear distribution of GFP-tagged WT or Y894AS897A-mutated Fragment A (Mut2) when expressed in HEK293T cells alone or in combination with mScarlet-tagged PAS^737^. The %N indicates quantification of % nuclear GFP localization.

Next, we sought to identify the PIM-binding region in the N-terminal PAS^737^ fragment. First, we generated truncations in this region (Figure 3A) and measured their interactions with Fragment A (841-1323). Our results indicated that the first 307 amino acids (termed PAS^307^), which contain only the PAS-A domain and unstructured flanking regions, are necessary to interact with Fragment A (Figure 3B). While examining multiple sequence alignments of the PIM region and PAS-A domain, we noticed the occurrence of reciprocal substitutions in PASK across different species (Figure S2). For instance, while S897 of PIM is conserved across most species, *G. gallus, D. melanogaster,* and *A. gambiae* PASK contain asparagine (N) or glutamate (E) at the equivalent position, respectively (Figure S2, red rectangles). The PAS-A region contains glutamate at 223 position of the PAS-A domain in most species; however, the aforementioned *G. gallus, D. melanogaster,* and *A. gambiae* PASKs contain lysine (*G. gallus*), serine (*D. melanogaster*), or threonine (*A. gambiae*) (Figure S2, red rectangles). Thus, it appears that the reciprocal changes in PIM and PAS-A domain at the human PASK 223 and 897 positions might preserve the interactions between these regions throughout evolution. Furthermore, K132 and W216 are two highly conserved amino acids in the PAS-A domains of PASK across invertebrates and vertebrates. To test if these residues contribute to PAS-A:PIM interactions, we mutated K132, E223, and W216 in the human PAS^737^ (K132A/W216A/E223A), which abrogated PAS-PIM interaction in a co-immunoprecipitation experiment from cells (Figure 3C). These results revealed the PAS-A domain of PASK to be engaged in a novel short linear interacting motif (SLIM) mediated interaction, which could be targeted for regulation. Finally, we asked if mutations in PAS-A that disrupted its interaction with PIM in cells (Figure 3D) affect the nuclear accumulation of PASK. We expressed GFP-tagged Fragment A and mScarlet-tagged WT or K132A/W216A/E223A-mutated PAS^737^ (PAS^737-Mut^) Fragments. As shown in Figure 3D-E, when co-expressed with mutated PAS^737^, Fragment A showed increased nuclear levels compared with WT-PAS^737^.

**Figure 3:**
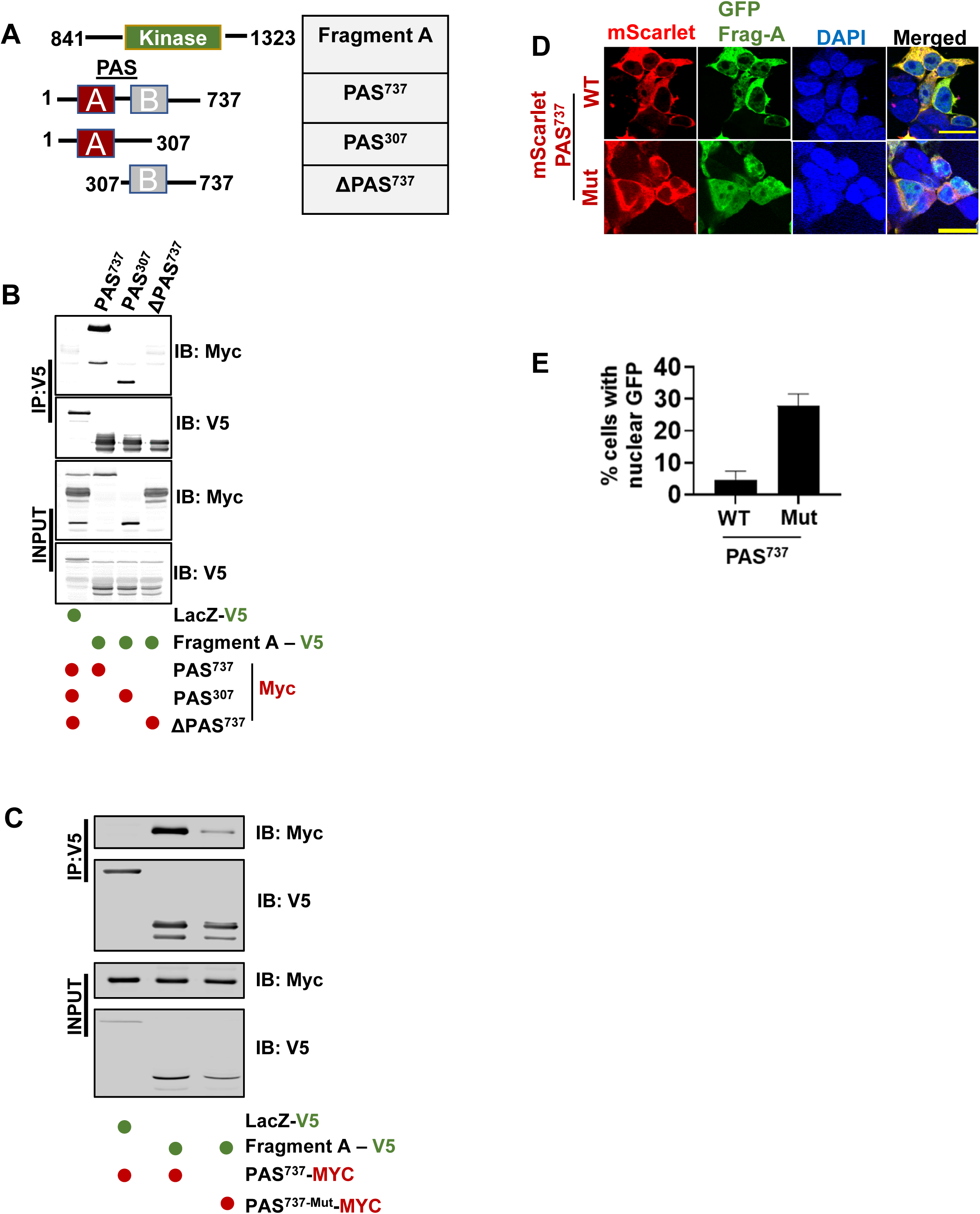
Mapping N-terminal residues in PAS-A domain that interact with PIM sequence. **(A)** Domain truncation analysis of PAS^737^ region that interacts with Fragment A. **(B)** Various fragments of PAS^737^-Myc were expressed in HEK293T cells and co-immunoprecipitation with V5-LacZ (negative control, lane 1) or Fragment A was conducted as described in Materials and Methods. **(C)** Myc-tagged WT or Mut (PAS^737-Mut^, K132A/W216A/E223A) PAS^737^ was co-expressed with V5-tagged Fragment A (lanes 2 and 3) in HEK293T cells. Interaction between proteins was determined by co-immunoprecipitation and western blot using the indicated antibodies as identified in the Figure panel. **(D)** Relative nuclear distribution of GFP-tagged Fragment A when co-expressed with either mScarlet-PAS^737^-WT or mScarlet-PAS^737-Mut^ (K132A/W216A/E223) **(E)** Quantification of images in Figure 3D showing % cells with nuclear GFP expression when co-expressed with either mScarlet-PAS^737^-WT or mScarlet-PAS^737^-Mut (K132A/W216A/E223A).

### Characterization of PAS-A/PIM interactions *in vitro*

We previously solved a solution structure of the PASK PAS-A (131-237) domain, identifying that small molecules can bind into a cavity inside of it as part of this work [9]. To confirm the PAS-PIM interaction in solution and determine how that might affect this PAS-A cavity, we first measured the binding affinity (K_D_) of PIM (AA 879-904) for the isolated PAS-A (131-237) domain using protein-detected NMR titrations [10]. By selectively monitoring PIM-induced chemical shift changes within uniformly ^15^N-labeled PAS-A with ^1^H/^15^N HSQC spectra across increasing concentrations of PIM (0.1– 2.5 mM), we measured an apparent K_D_ of 2.4 mM of PIM for PAS-A (Figure 4A). To identify the residues perturbed by PAS-A:PIM interactions, we compared changes in backbone amide ^1^H/^15^N shifts from unliganded PAS-A upon the addition of either PIM or the KG-571 small molecule, which we previously demonstrated to bind inside of the PAS-A cavity (compound 1 in [9]) (Figures 4B, C). These analyses showed chemical shift changes by either PIM or KG-571 around the PAS-A cavity, including perturbations in the Aβ strand, Fα helix, FG-loop, and Gβ, Hβ, Iβ strands [9]. We note that these changes were in complementary groups of residues in the PAS-A solution structure: PIM affected sites adjacent to the cavity, with the largest changes in the FG loop and flanking regions, while KG-571 bound within the cavity consistent with our prior observations [9]. Interestingly, we observed substantial chemical shift perturbations for W216 (Hβ), and E223 (HI-loop) upon the addition of PIM and KG-571, while K132 (Aβ) was perturbed only with KG-571, but not PIM, possibly due to the missing N-terminal residues within our NMR constructs which may facilitate transient PAS-A:PIM interactions in the cell-based assay with the longer PAS^737^ fragment.

**Figure 4:**
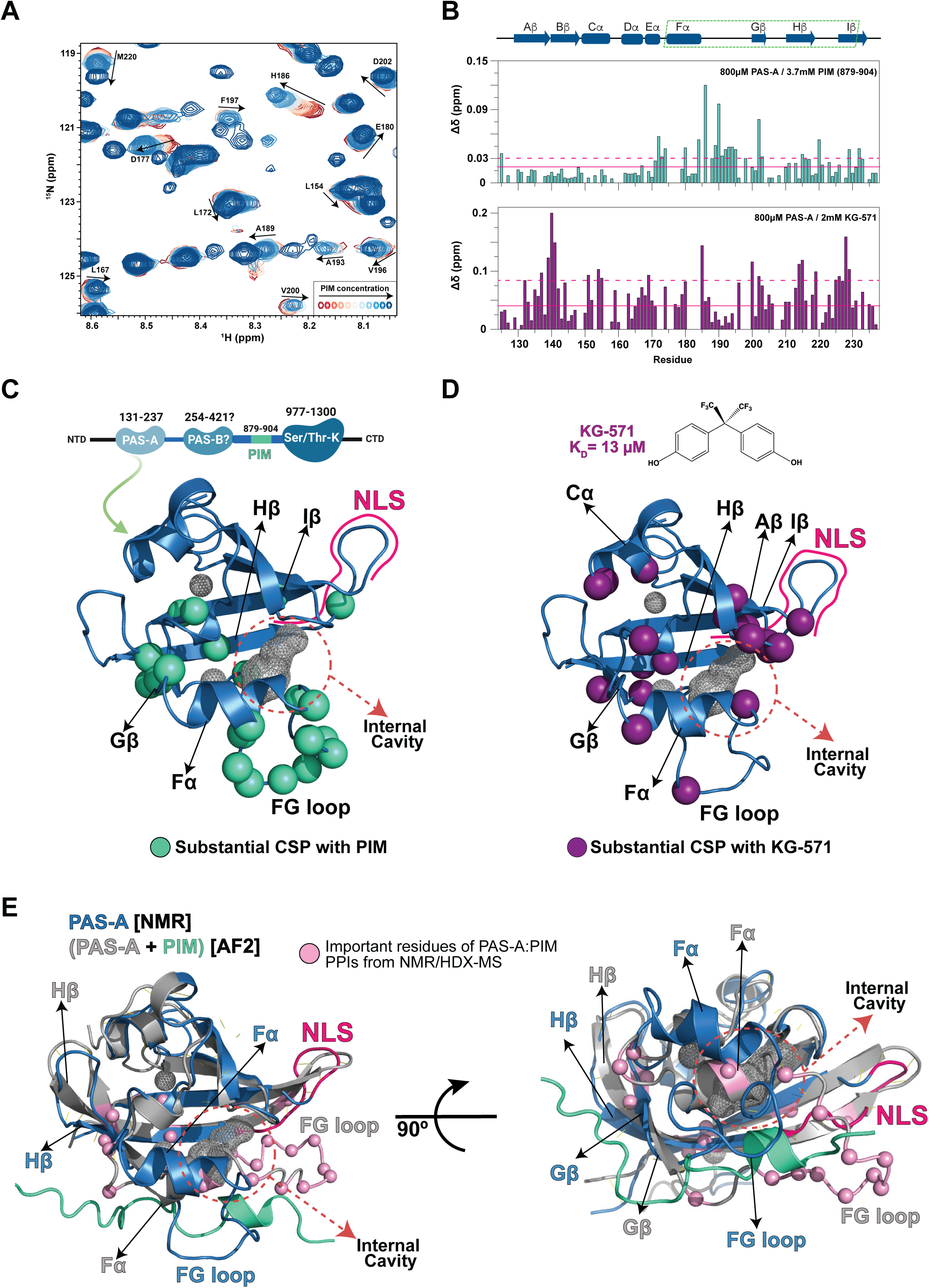
NMR-based identification of PIM binding site on PAS-A. **(A)** ^1^H/^15^N HSQC spectra of ^15^N-labeled PAS-A (131-237) in the presence of 0.1-2.5 mM unlabeled PIM (879-904), showing ligand-dependent chemical shift perturbations (CSPs) indicating binding. **(B)** Residues perturbed by the addition of either PIM (top) or KG-571, a previously-described small molecule ligand (compound 1 in [9]) (bottom), as identified by CSPs in backbone amide ^1^H/^15^N chemical shift changes from apo PAS-A. Changes were quantified by using Δδ(ppm)=(Δδ ^2^+(0.2*Δδ)^2^)^0.5^; solid red lines indicate the average level of observed CSPs, while dashed lines indicate the average + one standard deviation of CSPs. **(C)** Solution structure of PAS-A (PDB-ID: 1LL8 [9]) showing the locations of residues with substantial CSPs (> average + one standard deviation) by the addition of 3.7 mM PIM. **(D)** Solution structure of PAS-A showing that the addition of 2 mM KG-571 induces substantial CSPs (> average plus one standard deviation) in residues which cluster around the internal cavity identified in the structure. **E)** An overlay of the experimental PAS-A structure (blue, PDB-ID: 1LL8 [9]) and an AlphaFold2 Multimers [28] model of PAS-A(131-237) and PIM(879-904) (grey and aqua, respectively), suggesting PIM interacts with PAS-A via regions adjacent to the internal cavity, including portions of the Fα helix, FG loop, Gβ strand, and Hβ strand. PAS-A residues showing substantial PIM-induced changes in NMR/HDX-MS experiments are shown as pink spheres.

To further characterize the structural rearrangements caused by PIM or KG-571 binding to PAS-A, we also used hydrogen-deuterium exchange monitored by mass spectrometry (HDX-MS), which reports on changes in backbone amide protection from exchange with solvent deuterons. We used local HDX-MS, where deuterium uptake is monitored at the peptide level to yield information on structural dynamics with more dynamic solvent-accessible areas more readily exchanging with deuterons than less solvent-accessible areas [11]. Similar to our NMR results, we observed higher deuterium exchange around the PAS-A cavity for the apo state compared to those with either PIM or KG-571 added (Figure S3). With PIM or KG-571, we observed less deuterium exchange (= stabilization) around the Cα, Dα, Fα helices, and FG-loop; we also observed PIM-specific in the Aβ strand and KG-571-specific stabilization in the Hβ-Iβ strands (Figure S3). We conclude from these results that structural changes within the hydrophobic core in or around the cavity of the PAS-A domain by PIM or ligand binding lead to changes in the structure and/or dynamics of regions of PAS-A involved in intramolecular interactions with other parts of PASK, giving rise to its functional regulation [9].

To visualize this interaction, we generated an AlphaFold2 (AF2) Multimers [27, 28] model of the PAS-A and PIM complex. While this model was generated without any direct input from our experimental data, we note that it too suggests PIM binds near the PAS-A cavity (Figure 4E). The model of the PAS-A:PIM complex with the best pLDDT score suggests substantial conformational changes occur upon PIM peptide binding, especially in the FG-loop, Fl1J, Gβ, Hβ regions that showed similar changes in NMR and HDX-MS experiments. We analyzed the PAS-A:PIM interactions in the AF2 models with the HADDOCK PRODIGY server which uses contact-based prediction of protein-protein complexes [12, 13]. In this analysis, we observed that PIM residues Y894 and S897 contacted several points with PAS-A in the AF2 model, corroborating our prior co-immunoprecipitation data showing the importance of both residues for PIM interactions with PAS^737^ (Figure 2). Taken together, these data suggest a model where both ligand and PIM interact around the flexible PAS-A cavity, potentially facilitating allosteric control of inter-or intra-molecular interactions involved in PASK biological functions.

### Identification of A Regulatable Nuclear Localization Motif in the PASK PAS-A domain

Based on these *in cellulo* and *in vitro* studies, we considered the possibility that the disruption of an intramolecular PAS-A and PIM interaction could underline the signal-regulated nuclear trafficking of PASK. However, the mutation of PAS-A binding residues, Y894A/S897A in full-length PASK (PASK^PIM-Mut2^), did not induce nuclear translocation of full-length PASK (Figure 5A-B). This is not surprising since we have recently reported the identification of two strong Nuclear Export Sequences (NES) located between residues AA 401-410 (NES1) and AA 666-671 (NES2) in PASK (Figure 5C), the combination of which is powerful enough retain an engineered PASK in the cytosol even when fused to the strong SV40 NLS [8].

**Figure 5:**
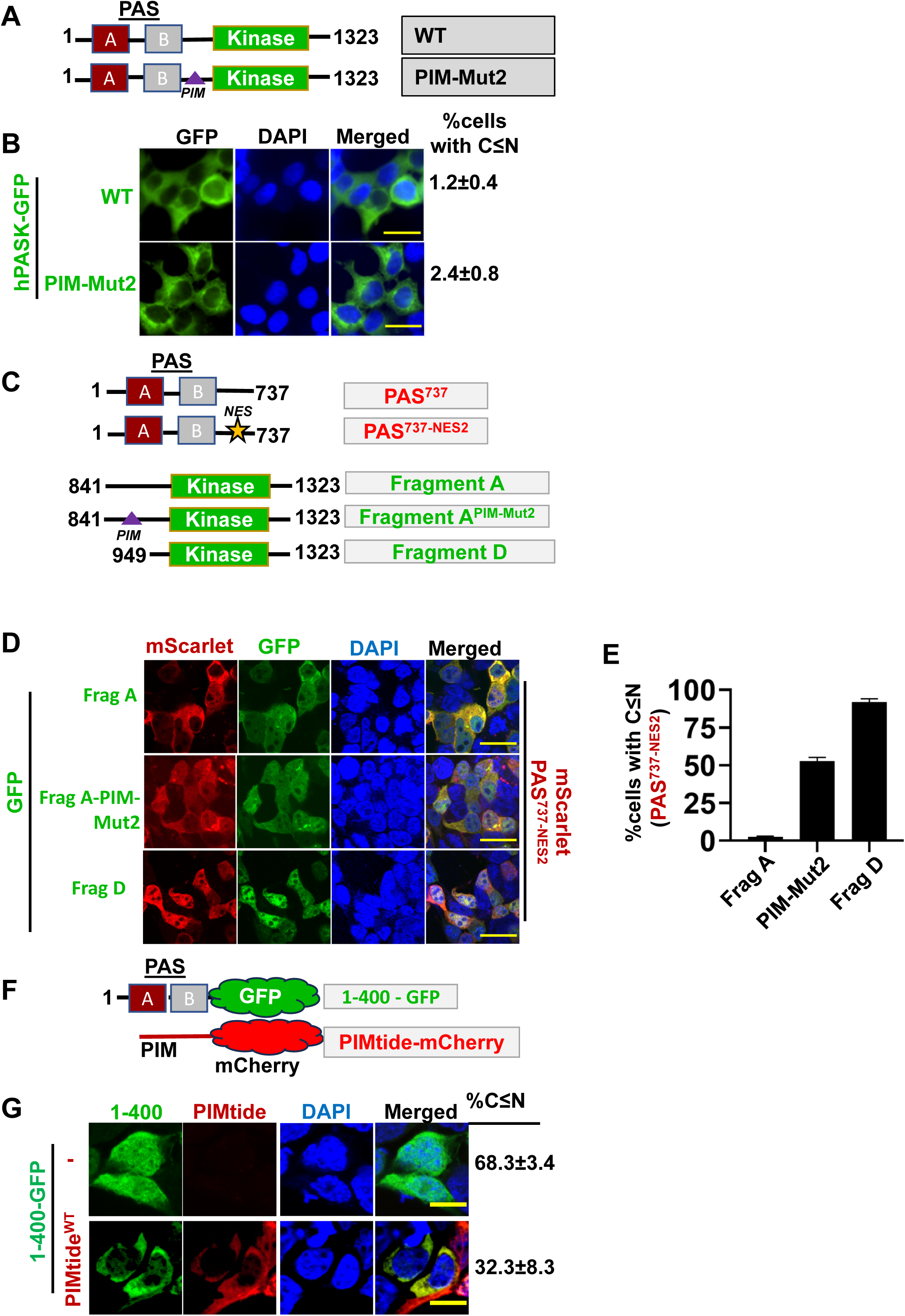
PIM interferes with the nuclear import of PAS-A. **(A)** Sequence and structural elements of WT or PIM-mutated (PIM-Mut2, Y894AS897A) full-length PASK used in Figure 5B. **(B)** GFP-tagged full-length WT or PIM-mutated (PIM-Mut2, Y894AS897A) full-length PASK were expressed in HEK293T cells, and their nuclear accumulation was observed by quantifying % cells with nuclear GFP (panel to the right). **(C)** Depiction of various N and C-terminal fragments used to identify the structural elements regulating PASK nuclear import. *NES=* mutations in nuclear export sequence (NES, L667SL669SL671S) **(D)** Immunofluorescence microscopy of mScarlet-PAS^737-NES2^ construct co-expressed with indicated GFP-tagged WT or PIM-Mut2 (Y894A/S897A) Fragment A or Fragment D. Scale bars = 20 µM. **(E)** Quantification of % cells with nuclear PAS^737^ expression when co-expressed with WT or Mut2-Fragment A or Fragment D. **(F)** Schematic representation of constructs used in Figure 5G. **(G)** GFP-400 (hPASK AA 1-400 fused to GFP) was expressed with or without PIMtide (AA 879-904)-mCherry in HEK293T cells. The GFP nuclear accumulation was observed by quantifying % cells with nuclear GFP (panel to the right). Scale bars = 20 µM.

To further understand the functional interplay of the PAS-A:PIM interaction and these nuclear export sequences, we co-expressed PAS^737^ or NES2-mutated version of PAS^737^ fragment (PAS^737-NES2^, PASK 1-737 with L667S/L669S/L671S mutations) (Table 1) along with WT or PIM-Mut2 Fragment A (PASK AA 841-1323 with Y894A/S897A) (Figure 5C-D). As expected, PAS^737^ was cytoplasmic regardless of the co-expression with either WT or a Y894A/S897S mutated Fragment A, owing to the presence of functional nuclear export sequences. Furthermore, PAS^737-NES2^ was retained in the cytoplasm when co-expressed with WT-Fragment A. However, co-expression of PAS^737-NES2^ with PIM-Mut2-Fragment A, which likely disrupted the interaction between PAS-A:PIM, resulted in an increased distribution of PAS^NES2^ construct in the nucleus, along with PIM-mutated Fragment A (Figure 5D-E).

We previously showed that a smaller PAS domain-containing fragment, Fragment 1-400 is predominantly nuclear since it lacks both nuclear export sequences in PASK but retains nuclear import capability [8]. To determine if interaction with PIM affects the nuclear import of the 1-400 fragment, we expressed GFP-tagged 1-400 fragment (Table 1, termed GFP-400) with or without mCherry-tagged PIMtide^WT^ (mCherry-879-904). Interestingly, co-expression with PIMtide^WT^ reduced the nuclear localization of GFP-400 (Figure 5F-G). These observations raised an intriguing possibility that the interaction between PAS-A and PIM controls the nucleo-cytoplasmic distribution of PASK and that PAS-A and flanking regions can mediate the import of PASK into the nucleus when it is not masked by PIM.

To directly test this hypothesis, we co-expressed mScarlet-tagged PAS^737^ or PAS^737-NES2^ proteins with GFP-tagged PIMtide^WT^ (GFP-AA 879-904) or PIMtide^Mut2^ (GFP-AA 879-904^Y894AS897A^) plasmids (Table 1, Figure 2C). We expected PAS^737^ to be excluded from the nucleus under all conditions due to the presence of two functional nuclear export sequences (Figure 6A-B). In contrast, PAS^737-NES2^ would be excluded from the nucleus if its nuclear localization motif was masked by PIM interaction. Interestingly, while nuclear import and accumulation of PAS^737-NES2^ were blocked when co-expressed with GFP-PIMtide^WT^, we noticed markedly increased nuclear import and retention of PAS^737-NES2^ when only when interaction deficient GFP-PIMtide^Mut2^ was co-expressed. These results show that the PIM occupancy in the PAS domain blocks nuclear import pathways (Figure 6B).

**Figure 6:**
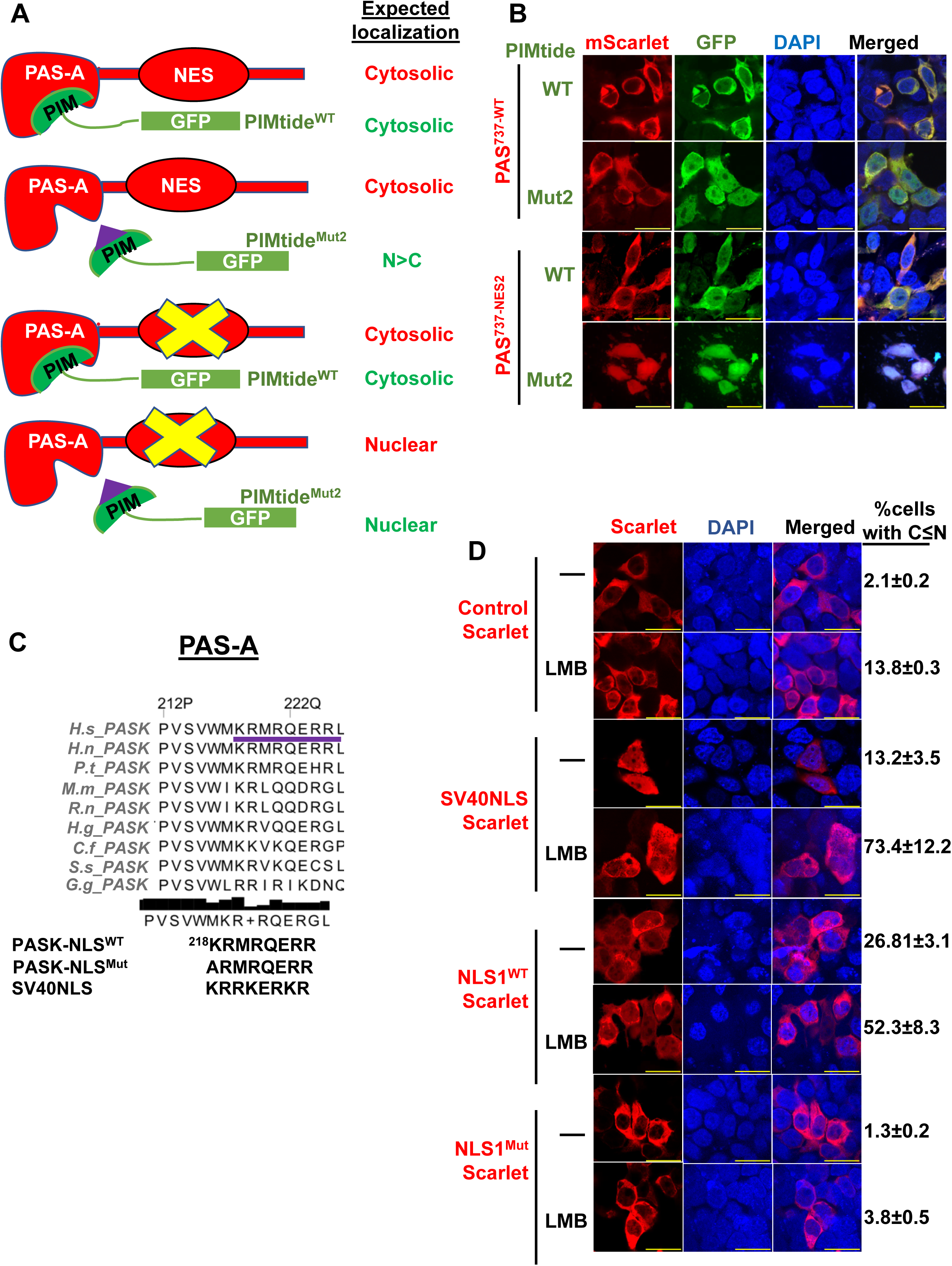
PIM masks a putative nuclear localization motif in PAS-A. **(A)** Proposed model describing the relationship between PIM binding and PAS-A domain nuclear import and expected outcome in a co-expression study. **(B)** mScarlet-tagged PAS^737-WT^ or PAS^737-NES2^ (PASK AA 1-^737(L667SL669SL671S)^) fragments were co-expressed with WT or Y894A/S897A (PIMtide^Mut2^) mutated GFP-tagged PIMtide. 24 hr after transfection, the relative nuclear levels of mScarlet-PAS^737^ fragments were visualized by confocal microscopy. Scale bars for all panels = 40µM. **(C)** The location of putative NLS residues in the PAS-A domain of PASK (underlined in red). **(D)** LacZ-mScarlet fusion protein with or without N-terminal NLS from SV40 (PKKKRKR), WT (KRMRQERR) or mutated PASK NLS (ARMRQERR) was expressed in HEK293T cells. 24 hr after transfection, cells were treated with 25nM leptomycin B for 2 hours. Relative nuclear levels of various constructs are quantified as % cells with nuclear mScarlet onto the right. Scale bars = 40µM.

Based on our analyses, we hypothesized that the PIM binding to the PAS-A domain masks a <u>N</u>uclear <u>L</u>ocalization <u>S</u>equence (NLS). To test this hypothesis, we first sought to identify the nuclear localization sequence in the PAS-A domain of PASK. Several bioinformatic analysis algorithms predicted residues K218-R225 (KRMRQERR) as a putative monopartite nuclear localization motif within the PAS-A domain (Figure 6C). Additionally, we also observed the segment of PAS-A HI-loop, which includes the NLS sequence (K218-R225), to have chemical shift changes in NMR with KG-571 and PIM (Figure 4B).

To directly test if residues K218-R225 function as NLS and are sufficient to import a heterologous protein, we generated a LacZ-mScarlet fusion protein (MW = ∼100kDa) with a peptide consisting of AA K218-R225 (^218^KRMRQERR^225^) from PAS-A domain (termed NLS1^WT^), or single point mutant version, NLS1^mut^ (^218^<u>A</u>RMRQERR^225^). As a positive control, we fused the SV40NLS (PKRKRRR) peptide, which is known to import tagged protein into the cell nucleus (14) with LacZ-mScarlet protein, and nuclear accumulation of LacZ-mScarlet alone was used as a baseline. We tested the ability of these constructs to mediate the nuclear import of LacZ-mScarlet fusion protein. We also used a nuclear export inhibitor, leptomycin B (LMB), to ensure nuclear retention of the imported fusion protein. As shown in Figure 6D, peptide NLS from SV40 or PAS-A domain of PASK (K218-R225) were comparable at stimulating the nuclear import of LacZ-mScarlet fusion protein in LMB-treated cells. On the other hand, K218A-mutated NLS1 (NLS1^mut^), was ineffective at promoting the nuclear localization of the LacZ-mScarlet fusion protein (Figure 6D). Thus, our results revealed a unique functional role of the intramolecular interaction between the PAS-A domain and PIM sequence in regulating the PASK nuclear translocation by masking the NLS sequence in the PAS domain.

### Metabolic regulation of PASK nuclear import via unmasking of PAS-PIM interaction

PASK is important for the terminal differentiation of embryonic and adult stem cells. While PASK is a predominantly cytoplasmic protein in proliferating stem cells, it is nuclear localized at the onset of myoblast differentiation. This nuclear distribution of PASK is induced by differentiation signaling cues and drives its physical association with its nuclear substrate, Wdr5. In addition, the nuclear export inhibitor, LMB, is ineffective in blocking PASK nuclear export under steady-state conditions. However, under serum-stimulated conditions, LMB effectively blocked the nuclear export of PASK, indicating that the nuclear import of PASK is signal-regulated. Mitochondrial glutamine metabolism induces CBP/p300-induced acetylation of PASK, which drives its nuclear localization [8]. Based on these results, we hypothesized that signaling pathways target PAS-PIM interactions to stimulate its nuclear import at the onset of terminal differentiation. Consistent with this hypothesis, we find that serum starvation resulted in cytoplasmic retention of PAS^737^ fragment when co-expressed with WT-Fragment A. Acute serum stimulation, on the other hand, induced strong nuclear localization of both PAS^737^ fragment and Fragment A (Figure 7A-B). Finally, since mitochondrial glutamine metabolism drives PASK nuclear translocation, we asked if glutamine metabolism serves to unmask the PAS-PIM interaction to stimulate PASK nuclear translocation. For this experiment, we used PAS^737-NES2^ co-expression with WT-Fragment A, both of which show cytoplasmic localization (see Figure 5D) under steady-state conditions. The use of PAS^737-NES2^ allowed for nuclear accumulation of the PAS^737^ fragment and circumvented the need for using LMB pretreatment. As shown in Figure 7C-D, in glutamine-starved cells, the PAS^737-NES2^ fragment remained cytosolic when co-expressed with WT-Fragment A. However, acute glutamine stimulation alone (in the absence of serum) was sufficient to increase the nuclear import of PAS^737-NES2^ and Fragment A.

**Figure 7:**
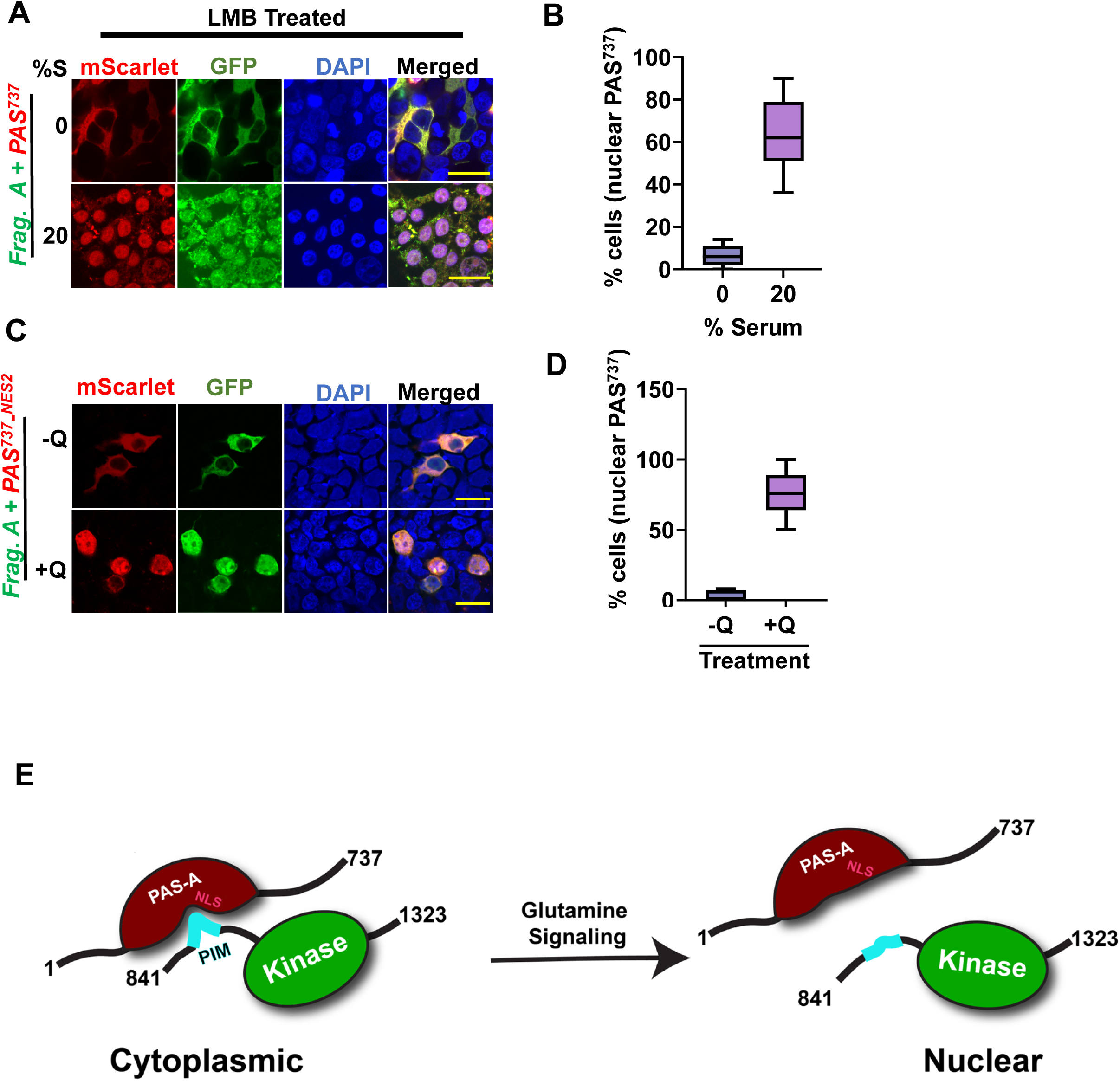
Signal-regulated co-import of N and C-terminal domains of PASK. **(A)** mScarlet-PAS^737^ and GFP-Fragment A were co-expressed in HEK293T cells. Cells were serum-starved for 18 hr. Cells were either refed with serum-free media (0%) or stimulated with 20% serum for 2 hr in the presence of 10nM of nuclear export inhibitor, LMB. Cells were fixed, and subcellular distribution was visualized by confocal microscopy. Scale bars = 40 µM. **(B)** Quantification of % cells with nuclear PAS^737^ from three independent experiments in Figure 7A. **(C)** mScarlet-PAS^NES^ and GFP-Fragment A were co-expressed in HEK293T cells. Cells were serum and glutamine (Q) starved for 18 hr. Cells were either refed with serum and glutamine-free media (-Q) or stimulated with 2mM glutamine alone for 2 hr. Cells were fixed, and subcellular distribution was visualized by confocal microscopy. Scale bars = 40 µM. **(D)** Quantification of % cells with nuclear PAS^737^ from three independent experiments in Figure 7A. **(E)** Model depicting possible mechanism for signaling control of PASK nuclear import mediated by glutamine metabolism that impinges on PAS-PIM interaction. While data here is based on *in trans* interactions between separated PAS^737^ and PIM-kinase Fragments, as shown in the Figure, we suggest that the same interactions occur *in cis* within full-length PASK.

Taken together, we reveal a novel metabolic regulation of PASK nuclear import via disruption of the intramolecular interaction between the N-terminal PAS domain and the C-terminal PAS interacting motif (PIM) (Figure 7E). Our results discovered a functional role of the PAS domain of PASK in mediating its nuclear import, which could play a crucial role in regulating the transition between self-renewal and differentiation of stem cells.

## Discussion

### Intramolecular interaction involving the PAS-A domain as a regulatory feature of the PASK function

The modes by which PAS domains sense and regulate cellular metabolic state are as diverse as the family members it comprises [1–4]. In this study, we identify a novel mode by which the PAS domain of PASK regulates the subcellular distribution of PASK in response to a metabolic cue. Our studies identified a protein-protein interaction interface in the PASK PAS-A domain that is intramolecularly occupied by a short-linear motif, which we termed PIM, situated adjacent to the C-terminal kinase domain. Through combined *in situ* biochemical and cell biological studies, we revealed the presence of a putative NLS in the PAS-A domain of PASK. Residues surrounding the PAS-A NLS interact with PIM, and mutations in PIM that disrupt this binding promote nuclear retention of an N-terminal PASK fragment which includes the PAS-A domain. A metabolic process likely regulates the dynamic of the PAS-A:PIM interaction since acute glutamine stimulation results in the nuclear import of PASK in a PAS-A:PIM interaction-dependent manner. Thus, our study identified the intramolecular association of PASK as a regulatory feature that could be controlled by metabolic pathways (Figure 7).

### A mechanism for using PAS-A to control the nucleo-cytoplasmic distribution of PASK using cellular metabolite status

Our discovery of a NLS in the PASK PAS-A domain was surprising since NLS sequences are often associated with unstructured or coiled-coil regions connecting protein domains. However, several literature reports support the notion that intramolecular interactions involving PAS domains can play regulatory roles in controlling the nuclear translocation of PAS-containing proteins. For example, in plant phytochrome B, a NLS is situated in the C-terminal PAS-B domain and is unmasked in a light-dependent manner [14]. Similarly, NPAS4 and PERIOD exhibit control of nucleo-cytoplasmic distribution via intramolecular interaction involving PAS domains [15–17]. Within PASK, the NLS we identified (residues K218-R225) can independently function when removed the surrounding domain (Figure 6D); this is not surprising, as the isolated NLS peptide is likely in a random coil conformation. Within the PAS-A domain NMR structure [9], however, we see that this sequence is also in an extended conformation that is similar to the one adopted by the SV40 NLS when bound to importin alpha [18], without obvious steric clashes required for the whole PAS-A domain to bind the importin.

Notably, our solution NMR and HDX-MS data suggest that PIM binding to PAS-A masks the NLS, with residues in the Fα helix, FG loop, and Hβ strand reporting on this interaction. Our unassisted AF2 Multimers model – generated without using any of these data – strongly implies that PIM binding causes conformational changes in PAS-A that cause the FG loop to fold back upon the NLS region, blocking the required interactions with importins required for nuclear import. These observations from *in vitro* experiments on minimal fragments are consistent with the cellular data on larger systems (Figure 1-2).

Excitingly, our *in vitro* data show that these PAS-A:PIM interactions occur adjacent to the internal cavity that we previously identified to bind small molecule ligands [9], including the artificial KG-571 compound utilized here (Figure 4). This coincidence clearly sets up the potential for metabolite binding within the cavity to allosterically control PIM binding, thus, NLS exposure and PASK nuclear import (Figure 7E). While proof of this control requires the identification of a physiological ligand for this domain (or alternatively, higher affinity artificial ligands than those currently known), many other PAS domains are well known to regulate comparable PAS:protein interactions via changes in ligand occupancy or configuration [19, 20].

## Conclusion

Using complementary techniques of *in vivo* and *in vitro* protein-protein interaction and localization assays, we identified a novel functional role of PAS-A domain of PASK as a regulator of its nuclear import machinery. Our study suggests a model of N-terminal PAS domain protein/ligand and protein/protein interactions within PASK, building on our prior work demonstrating PAS-A could bind small organic compounds [9]. In this study we expand the analysis to show that the PAS-A domain engages in intramolecular interactions via a small linear motif, termed PIM that coordinates PASK nuclear import in accordance with metabolic inputs. We further identified specific residues and interaction sites between the PAS-A and PIM fragments, which could be key to the overall function of PASK. Taken together, our study could lead to further discovery of PASK biological regulation. As the nuclear import of PASK is critical for the stem cell differentiation program, our analyses of PAS-A: PIM interaction provide a significant mechanistic advance in understanding how metabolic signals drive PASK nuclear import at the onset of terminal differentiation program, which can be targeted to develop therapeutic strategies aimed at balancing stem cell self-renewal and differentiation capabilities.

## Acknowledgments

This work was supported by grants from the National Institutes of Health (R01 AR073906 to C.K.K., R01 GM106239 to K.H.G., F31 GM142258 to R.A.).

## Conflict of Interest Disclosure

C.K.K. is an inventor on a provisional U.S. patent application covering the potential uses of PIM peptides as modulators of stem cell differentiation.

## Materials and Methods

### Cell lines and cDNA transfections

HEK293T and C2C12 myoblasts were obtained from ATCC and cultured in standard Dulbecco’s Modified Eagle’s Medium (DMEM) with 10% Fetal Bovine Serum, with glutamine and sodium pyruvate (Gibco, Cat# 119995) and 1% penicillin and streptomycin. For protein expression, cells were transfected with plasmids as indicated the figure legends using polyethylenimine (PEI) based transfection.

### Cloning and mutagenesis

PCR-based amplification was used to generate C-terminally truncated domains in pCDNA3.1A-V5 vector. Similarly, N-terminal fragments were cloned into pCDNA3.1B-Myc vector following PCR-based amplification. For site-directed mutagenesis, sewing PCR was used using partially overlapping primers with a centrally located mutant nucleotide [21].

### Cell Lysis, Protein Extraction and Co-Immunoprecipitation

The protein lysates from overnight transfected HEK293T cells when the cells reached approximately 80% confluency. For that, cells were harvested by scraping and subsequently lysed in native cell lysis buffer. The native cell lysis buffer consisted of 20 mM Tris-HCl pH 7.5, 150 mM NaCl, 1 mM EDTA, 1 mM NaF, 1 mM beta-glycerophosphate, and 1% Triton X-100, supplemented with freshly added protease and phosphatase inhibitors including 1 mM PMSF, 1× protease inhibitor cocktail, and 1× phosphatase inhibitor cocktail (Sigma). The cell lysates were incubated on ice for 20 min to ensure complete lysis, followed by centrifugation at 15K RPM X 10mins in refrigerated centrifuge.

For co-immunoprecipitation, the clarified cell lysates were utilized as described above. Ten percent of each lysate was collected as input, and the remaining lysate was subjected to immunoprecipitation (IP) overnight at 4°C using either Anti-V5 Agarose Affinity Gel beads (Sigma, A7345) or Anti-FLAG M2 magnetic beads (Sigma, M8823), depending on the target protein. The IP samples were then washed five times on ice using either wash buffer (20 mM Tris-HCl pH 7.5, 150 mM NaCl, 1% Triton X-100). Subsequently, the samples were resolved by SDS-PAGE, followed by immunoblot analysis using the specific antibodies as indicated in figure panels.

### Immunofluorescence microscopy and data quantification

For immunofluorescence microscopy, cells were seeded in 24-well plates on glass coverslips. The coverslips were precoated with 0.1% gelatin for HEK293T or C2C12 cells. At the designated experimental time points, the cells were fixed with 4% paraformaldehyde (PFA) in 1× PBS for 15 min, followed by three washes with 1× PBS. To visualize fluorescently tagged proteins, coverslips were mounted using ProLong Antifade mounting media containing DAPI (ThermoFisher Sci) and confocal analysis was conducted after mounting media curing. For detection of non-fluorescently labeled proteins, cells were permeabilized using 0.2% Triton X-100 in 1× PBS for 10 min at room temperature. Subsequently, blocking was performed using 10% normal goat serum in 1× PBS for 1 hour. Primary antibodies, diluted in 1× PBS, were added to all wells and incubated overnight at 4°C in a humidified chamber. Afterward, the wells were washed three times with 1× PBS. For secondary antibody staining, appropriate secondary antibodies (Alexa Fluor 488 – green and 568 – red, Thermo Fisher Sci) were added to the wells and incubated for 1 hour at room temperature. Following three washes with 1× PBS, the coverslips were mounted onto glass slides using ProLong Antifade mounting media with DAPI (Thermo Fisher Sci). The mounted coverslips were allowed to cure overnight at room temperature before imaging was performed using confocal microscopy (Nikon A1R). Analysis and quantification of the acquired images were carried out using calibrated Fiji software [22].

### Protein Expression and Purification

The hPASK PAS-A domain (residues 131-237) was expressed as an N-terminal His_6_-tagged fusion [23] in *E. coli* BL21(DE3) using ^15^NH_4_Cl-containing M9 minimal media as previously described [9]. Harvested pellets from 1 L growths were suspended in 40 mL 50 mM Tris-HCl (pH 6.5), 100 mM NaCl, and 25 mM imidazole buffer and sonicated at 4 °C. Lysates were separated by centrifugation at 77500 x g at 4 °C for 45 min, with the resulting supernatants filtered with 0.2-micron filters and subjected to Ni-Sepharose affinity purification, with the pHis-PAS-A protein obtained by gradient elution with 25–500 mM imidazole in the same buffer. Eluted samples were digested with His_6_-TEV protease [24] overnight to specifically cut a TEV protease site present between the His_6_ affinity tag and the PAS-A domain, with the digesting sample dialyzed into 3 L of imidazole-free buffer at 4 °C. The cleaved protein was again applied to Ni-Sepharose affinity chromatography, this time collecting the PAS-A protein in the flow-through fraction. Fractions were concentrated (Amicon Ultra, Millipore) and subjected to a final Superdex 75 size exclusion chromatography step in 20 mM sodium phosphate (pH 6.5), 50 mM NaCl, 5 mM DTT, and 6 mM NaN_3_. Fractions corresponding to monomeric PAS-A were concentrated, and flash frozen in liquid N_2_ and stored at −80 °C. The purity of the protein was assessed via SDS-PAGE electrophoresis, with typical yields of 100-200 mg/L.

### NMR Data Acquisition and Analysis

All NMR experiments were conducted at 298 K on Bruker Avance III HD NMR spectrometers at 700 or 800 MHz equipped with 5-mm inverse TCI cryoprobes, pulsed-field Z (700 MHz) or XYZ (800 MHz) gradients, and Topspin 3.5 software (Karlsruhe, Germany). NMR experiments of PAS-A (0.1 mM) and PIM (residues 879-902) titration (0.1, 0.3, 0.5, 0.7, 0.9, 1.2, 1.5, 1.7, 2.0, and 2.5 mM) were conducted in 14.7 mM Tris pH 7.4, 35 mM sodium phosphate (pH 7.4), 20 mM NaCl, and 10% D_2_O. The saturated concentration of artificial ligand KG-571 (2 mM) and PIM (3.7 mM) for NMR chemical shift perturbation data were collected with 225 µM of PAS-A in 20 mM sodium phosphate (pH 6.5), 50 mM NaCl, 5 mM DTT, and 6 mM NaN_3_. All NMR data were processed and analyzed with NMRFx and/or NMRViewJ (One Moon Scientific) [25, 26].

### HDX-MS Data Acquisition and Analysis

Protein samples for Hydrogen Deuterium Exchange Mass Spectrometry (HDX-MS) were prepared as described above in 20 mM sodium phosphate (pH 6.5), 50 mM NaCl, 5 mM DTT, and 6 mM NaN_3_. Stock protein concentrations were 100 µM PAS-A, with 2 mM KG-571 or 3 mM PIM added for complex samples (pH of the complexes were corrected for 6.5). Deuterium exchange was initiated at room temperature by diluting 5 µL of sample in 75 µL of D_2_O buffer, allowing exchange to continue for 30, 150, 300, 1000, 3000, and 5000 seconds before being quenched with 80 µL of ice-cold (∼ 3 °C) quench buffer (3 M GdHCl + 3% acetonitrile + 0.8% formic acid (pH ∼ 1.9)).

Quenched protein samples were immediately injected over an Enzymate BEH Pepsin Column (Waters) to digest the protein while desalting the resulting peptides by a 10 µL C8 Opti-lynx II trap cartridge (Optimize Technologies) at a flow rate of 0.12 mL/min using 0.25% formic acid in water as a mobile phase. After three min of injection, peptides were resolved with an analytical C18 column (Hypersil GOLD 1×50 mm, Thermo Scientific) and quickly eluted into a Bruker maXis-II ETD UHR ESI-QqTOF spectrometer at a flow rate of 40 μL/min for mass determination.

All steps of quenching, digestion, and injection were performed using a LEAP HDX automation robotic system (Trajan Scientific and Medical) with each exchange timepoint repeated twice for statistical validation. In addition to the HDX-MS samples noted above, unlabeled samples were run under identical conditions as HDX except in H_2_O buffer to provide masses and retention times of undeuterated peptides as reference. The unlabeled run was performed with MS/MS fragmentation to identify peptides using COMPASS DataAnalysis and BioTools software (Bruker). Every matched m/z and retention time pair was assigned to only one peptide sequence based on accurate mass and MS/MS measurements. This resulted in coverage of 100% of the protein sequence. The resulting peptide (sequence, m/z, charge state, and retention time) lists and the exchange rates were analyzed using HDExaminer version 3.3 (Sierra Analytics) software. All peptide matches were manually confirmed after automatic assignment by HDExaminer. At the end of the data analysis, ∼90% of the peptides resulted in high confidence coverage for the HDX-MS of the three reported conditions (apo PAS-A, PAS-A/PIM complex, PAS-A/KG-571 complex).

### AlphaFold2 Multimers Model Generation

To generate structural models of PAS-A:PIM complex, we used AlphaFold2 Multimers guided high-accuracy prediction [27, 28], the hPASK (Uniprot ID: Q96RG2) PAS-A(131-237) and PIM(879-904) sequences. The existing structure of PAS-A (PDB-ID: 1ll8) [9] was used as a template for the PAS-A:PIM complex model generation. Out of 10 models generated, the model shown in Figure 4E was the best among these as ranked by the pLDDT confidence measure with high accuracy. The model is available in ModelArchive (modelarchive.org) (ID: ma-96ezk [*available pre-publication with the accession code MvzBruyYQg*]).

## Notes

### Summary of Updates

Upon the peer review process, additional experiments were conducted, which resulted in a revised model of how nutrient signaling cues affect the intramolecular interactions in PASK to drive its nuclear import. We have also rearranged the figures and overall flow of the manuscript. Finally, substantial improvement was made in the quality of the microscopic images, clarification of various statistical analyses, and quantification.

