## Supplemental document for "Signal-regulated unmasking of the nuclear localization motif in the PAS domain regulates the nuclear translocation of PASK"

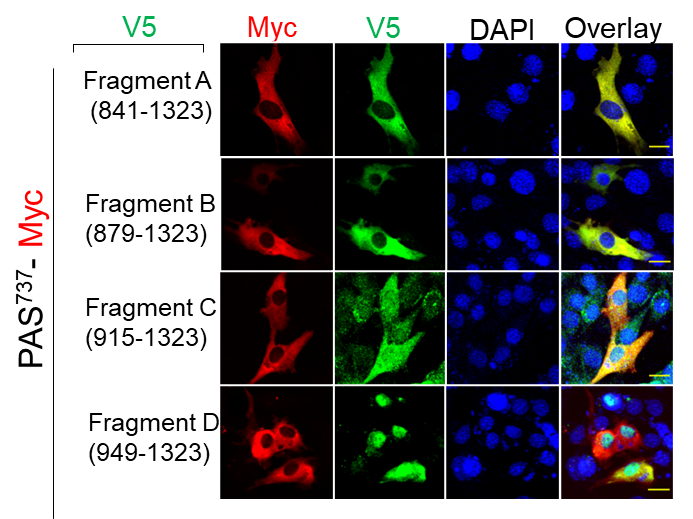

**Figure S1:** **Subcellular localization of C-terminal region of PASK in the presence of N-terminal domains.** C2C12 cells transfected with V5-tagged C-terminal fragments and Myc-tagged N-terminal PAS^737^ fragment. 24 hr after transfections, cells were fixed, and subcellular localization of V5 and Myc-tagged proteins was visualized by immunofluorescence microscopy as described in Materials and Methods.

PAS-A

PIM

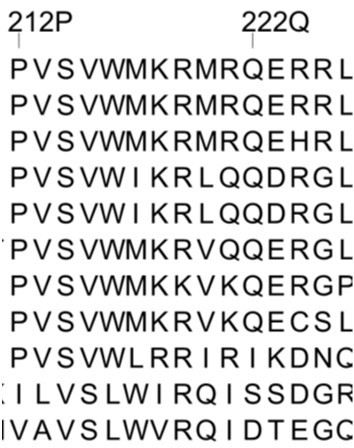

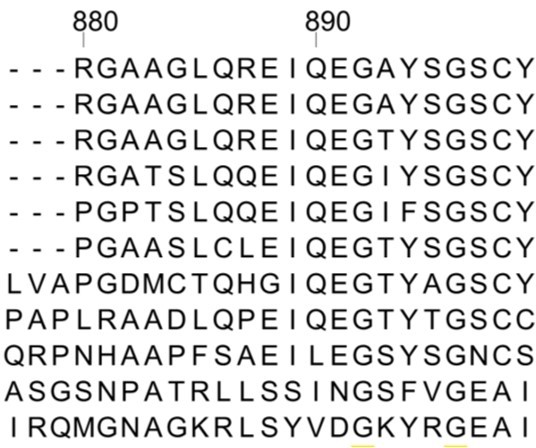

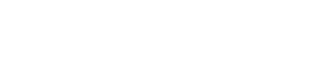

*H.s_PASK*

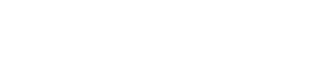

*H.n_PASK*

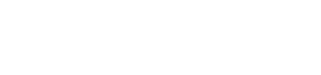

*P.t_PASK*

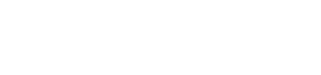

*M.m_PASK*

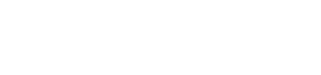

*R.n_PASK*

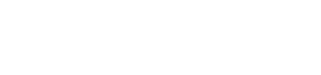

*H.g_PASK*

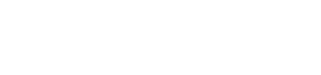

*C.f_PASK*

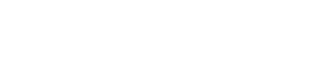

*S.s_PASK*

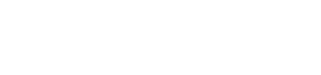

*G.g_PASK*

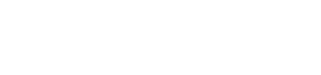

*D.m_PASK*

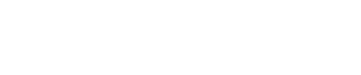

*An.g_PASK*

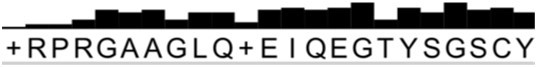

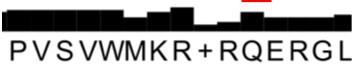

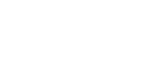

Y894

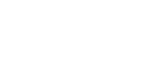

S897

**Figure S2: Multiple sequence alignment of C-terminal and N-terminal PAS-A domain of PASK.** Multiple sequence alignment was generated using Clustal Omega (1), and the conservation score was calculated using JalView. Red rectangular boxes indicate reciprocal substitution in the PIM and PAS-A binding region adjacent to the nuclear localization motif (see Figure 6).

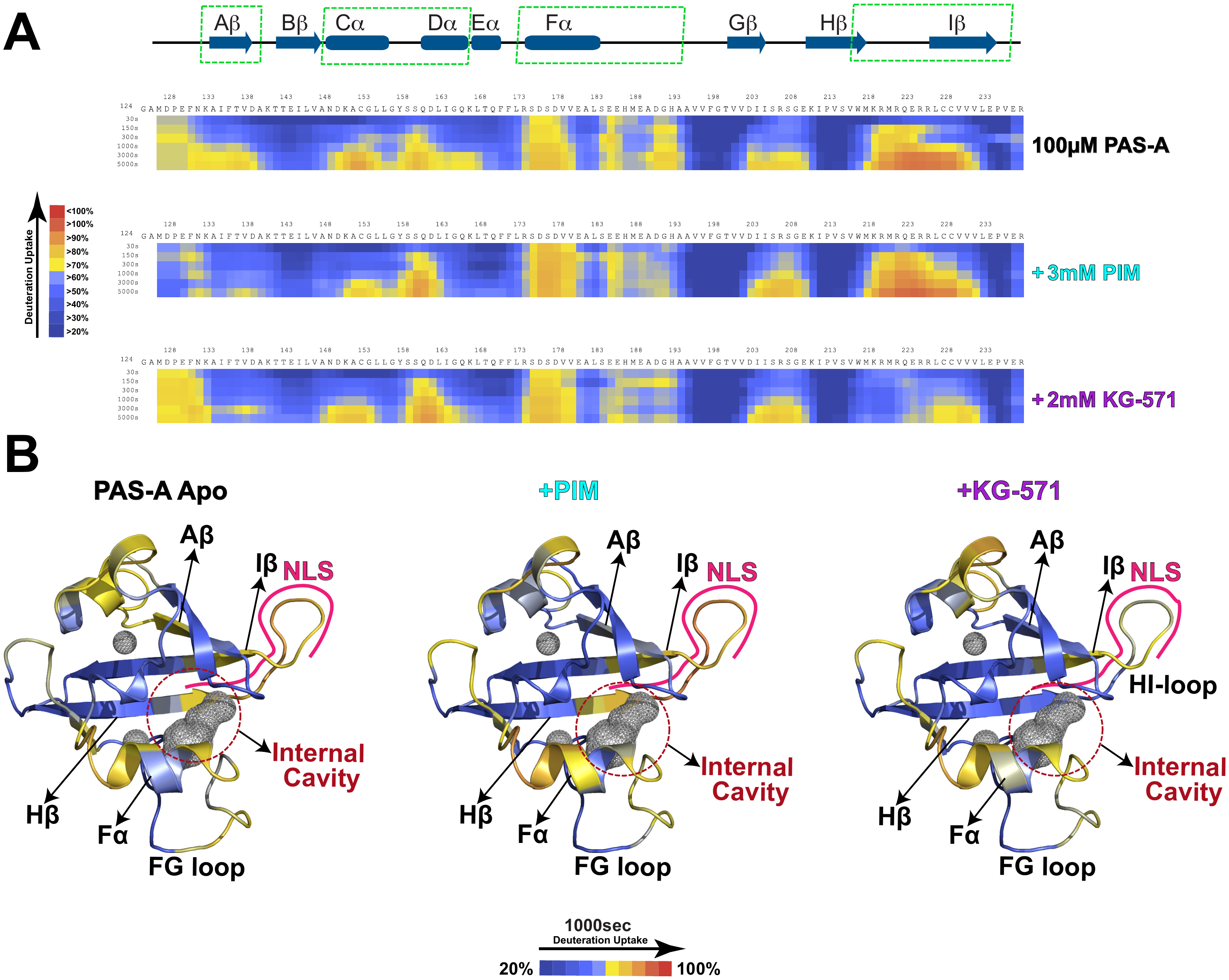

**Figure S3: HDX-MS comparison of overall structural dynamics of PAS-A with KG-571 and PIM interactions.** **A)** HDX-MS heatmaps indicating the progressive incorporation of deuterium as a function of incubation time of either PAS-A alone (top) or in the presence of either PIM (middle) or KG-571 (bottom) at indicated concentrations. Secondary structure from PAS-A solution structure (PDB-ID: 1LL8 (2)) schematically indicated above HDX-MS data, with green boxes highlighting areas with substantial change in HDX upon addition of either compound). **B)** Solution structure of PAS-A, with residues colored to indicate levels of deuterium incorporation at each amide site after 1000 seconds exchange under each of the three sample conditions indicated in panel A.
